# Contribution of *trans* regulatory eQTL to cryptic genetic variation in *C. elegans*

**DOI:** 10.1101/120147

**Authors:** L. Basten Snoek, Mark G. Sterken, Roel P. J. Bevers, Rita J. M. Volkers, Arjen van’t Hof, Rachel Brenchley, Joost A. G. Riksen, Andrew Cossins, Jan E. Kammenga

## Abstract

**Background:** Cryptic genetic variation (CGV) is the hidden genetic variation that can be unlocked by perturbing normal conditions. CGV can drive the emergence of novel complex phenotypes through changes in gene expression. Although our theoretical understanding of CGV has thoroughly increased over the past decade, insight into polymorphic gene expression regulation underlying CGV is scarce. Here we investigated the transcriptional architecture of CGV in response to rapid temperature changes in the nematode *Caenorhabditis elegans*. We analyzed regulatory variation in gene expression (and mapped eQTL) across the course of a heat stress and recovery response in a recombinant inbred population.

**Results:** We measured gene expression over three temperature treatments: i) control, ii) heat stress, and iii) recovery from heat stress. Compared to control, exposure to heat stress affected the transcription of 3305 genes, whereas 942 were affected in recovering animals. These affected genes were mainly involved in metabolism and reproduction. The gene expression pattern in recovering animals resembled both the control and the heat-stress treatment. We mapped eQTL using the genetic variation of the recombinant inbred population and detected 2626 genes with an eQTL in the heat-stress treatment, 1797 in the control, and 1880 in the recovery. The *cis*-eQTL were highly shared across treatments. A considerable fraction of the *trans*-eQTL (40-57%) mapped to 19 treatment specific *trans*-bands. In contrast to *cis*-eQTL, *trans*-eQTL were highly environment specific and thus cryptic. Approximately 67% of the *trans*-eQTL were only induced in a single treatment, with heat-stress showing the most unique *trans*-eQTL.

**Conclusions:** These results illustrate the highly dynamic pattern of CGV across three different environmental conditions that can be evoked by a stress response over a relatively short time-span (2 hours) and that CGV is mainly determined by response related *trans* regulatory eQTL.

## Background

Many organisms can respond to sudden changes in the ambient environmental conditions by adjusting their gene expression levels [1]. In particular, invertebrates are prone to environment-induced rapid gene-expression changes. For instance, gene expression in the nematode *Caenorhabditis elegans* can swiftly change due to exposure to pathogens, temperature, and toxicants [2–6]. A common denominator of many of these studies is that they provide snapshots in time of the responses elicited by these environments. As such, they provide static profiles of gene expression at a given moment. Further insight into the dynamics of gene expression and gene-expression regulation can be achieved by following responses, such as development or aging, over time [7–9].

Gene-expression regulation can be studied by investigating the transcriptional response in the context of natural variation using genetical genomics [10]. In this approach, a genetically segregated population (*e.g.* recombinant inbred lines, RILs) is used in a transcriptomics experiment to determine the genetic architecture of gene expression. Genetical genomics has been used in many species, including *C. elegans*, and many environments. One of the marked observations resulting from a comparison of different environments is the change in genetic architectures, revealing additional variation (as conceptualized by [11]). For example, the transcriptional architecture of *C. elegans* grown at 16°C or 24°C differs markedly [12]. Such studies show that the genetic architecture of gene expression is a dynamic process affected by relatively long-term differences in environment or age.

Gene expression is also highly dynamic during development and growth. For example, development in *C. elegans* is a tightly regulated process that has strongly correlated patterns of gene expression [8, 9, 13]. These developmental expression dynamics can be affected by natural genetic variation, for example between two commonly used divergent strains N2 and CB4856 [14]. Depending on the developmental stage, up to 10% of the genes in these strains show differential expression linked to genotype. This enabled the mapping of QTLs for gene expression dynamics during development [15].

Environmental changes can unlock genetic variation that remains hidden when in one condition but becomes apparent in another, a phenomenon called cryptic genetic variation (CGV) [16]. CGV has received quite some renewed interest over the past decade and it is suggested that CGV provides the raw material of evolution and adaptation under different environmental conditions [17]. Unlocking CGV can be achieved by altered gene-expression regulation, such as the transcriptional response to changing ambient conditions. Gene regulatory networks play an important role in understanding how environmental cues affect cryptic genetic variation [18]. Here we aim to investigate the CGV of the genetic architecture over the course of a strong environmental stimulus across a recombinant inbred line (RIL) population in the nematode *C. elegans*. As stimulus we used a heat stress, since *C. elegans* strongly reacts to temperature differences [19]. Furthermore, the two parental strains of the RIL population, Bristol (N2) and Hawaii (CB4856), display extensive variation in response to heat [12, 19–21]. First, we obtained the transcriptomes of the RILs over three treatments: (i) control, 48 hours at 20°C; (ii) heat stress, 46 hours at 20°C followed by 2 hours exposure to 35°C; (iii) recovery, same as the heat stress, followed by an additional 2 hours at 20°C. The transcriptomes were used for comparing the transcriptional differences as well as the identified eQTL between the three treatments. The treatments resulted in strong differences in gene expression, whereby the recovery treatment showed characteristics of both the control and the heat-stress treatment. Comparative analysis over the identified eQTL per treatment showed that *cis*-eQTL were strongly conserved over treatments. *Trans*-eQTL were more dynamic and display little overlap between treatments. We show that CGV is mainly manifested by *trans*-eQTL of specific sets of genes in specific environments. This makes the genetic architecture of gene expression variation an even more complex and cryptic phenomenon than previously thought.

## Methods

### Strains used

The wild-types N2 and CB4856 and 54 RILs derived from a CB4856 x N2 cross were used (strains generated in [12]). For 49/54 of these strains low-coverage sequencing was applied to construct a more detailed genetic map (see also [22]). A matrix with the strain names and the genetic map can be found in **Additional file 1**.

### Nematode culturing

The strains were kept on 6-cm Nematode Growth Medium (NGM) dishes containing *Escherichia coli* strain OP50 as food source [23]. Strains were kept in maintenance culture at 12°C, the standard growing temperature for experiments was 20°C. Fungal and bacterial infections were cleared by bleaching [23]. The strains were cleared of males prior to the experiments by selecting L2 larvae and placing them individually in a well in a 12-wells plate at 20°C. Thereafter, the populations were screened for male offspring after 3 days and only the 100% hermaphrodite populations were transferred to fresh 9-cm NGM dishes containing *E. coli* OP50 and grown until starved.

### Control, heat stress, and recovery from heat stress experiments for transcriptomics

The experiments were started by transferring a starved population to a fresh 9-cm NGM dish. This population was grown for 60 hours at 20°C to obtain egg-laying adults, which were bleached in order to synchronize the population. The eggs were transferred to a fresh 9-cm NGM dish. Three growing conditions were applied: (i) the control treatment was grown for 48 hours at 20°C, (ii) the heat-stress treatment was grown for 46 hours at 20°C followed by 2 hours at 35°C, and (iii) the recovery treatment was grown for 46 hours at 20°C, followed by 2 hours at 35°C and thereafter 2 hours at 20°C. Before the start of the treatment, the developmental stage of the population was determined by observing the developmental stage of the vulva in multiple individuals. Populations not consisting of L4 larvae were not isolated. Directly at the end of the treatment, the population was washed off the plate with M9 buffer and collected in an Eppendorf tube, which was flash frozen in liquid nitrogen. In this manner, 48 RILs per condition were assayed.

### Genotypes and genetic map construction

Previously, 49 lines were sequenced and aligned. The single-nucleotide polymorphism (SNP) calls per strain were taken for constructing the genetic map [22]. The SNP density was determined per 10 kb bins and recombination events were recognized as transition of an area where there were no CB4856 SNPs in 10 consecutive bins into an area where there were CB4856 SNPs and the other way around. It was not allowed to have two recombination events within 10 consecutive bins (100 kb). The 10 kb bin where the first SNPs were detected was marked as the recombination event. Before use in mapping, the map was filtered for informative markers – that is - markers indicating a recombination event in at least one of the lines. This resulted in a map of 729 informative markers, each indicating the location of the recombination events within 10 kb (see the figure in **Additional file 2**).

The genetic map was investigated by correlation analysis to assess the linkage between markers. Markers on the centers of the chromosomes showed strong linkage (see also [24]). No strong in between chromosome correlations were found (see the figure in **Additional file 3**).

### Transcript profiling

#### RNA isolation

The RNA of the RIL samples was isolated using the RNeasy Micro Kit from Qiagen (Hilden, Germany). The ‘Purification of Total RNA from Animal and Human Tissues’ protocol was followed, with a modified lysing procedure; frozen pellets were lysed in 150 µl RLT buffer, 295 µl RNAse-free water, 800 µg/ml proteinase K and 1% ß-mercaptoethanol. The suspension was incubated at 55°C at 1000 rpm in a Thermomixer (Eppendorf, Hamburg, Germany) for 30 minutes or until the sample was clear. After this step the manufacturer's protocol was followed.

#### cDNA synthesis, labelling and hybridization

The ‘Two-Color Microarray-Based Gene Expression Analysis; Low Input Quick Amp Labeling’ -protocol, version 6.0 from Agilent (Agilent Technologies, Santa Clara, CA, USA) was followed, starting from step five. The *C. elegans* (V2) Gene Expression Microarray 4X44K slides, manufactured by Agilent were used. Before starting cDNA synthesis, quality and quantity of the RNA were measured using the NanoDrop-1000 spectrophotometer (Thermo Scientific, Wilmington DE, USA) and RNA integrity was determined by agarose gel electrophoresis (3 μL of sample RNA on 1% agarose gel).

#### Data extraction and normalization

The microarrays were scanned by an Agilent High Resolution C Scanner with the recommended settings. The data was extracted with Agilent Feature Extraction Software (version 10.5), following manufacturers’ guidelines. Normalization of the data was executed in two parts, first the RILs and the ILs, second the mutant strains. For normalization, “R” (version 3.3.1 × 64) with the Limma package was used. The data was not background corrected before normalization (as recommended by [25]). Within-array normalization was done with the Loess method and between-array normalization was done with the Quantile method [26]. The obtained single channel normalized intensities were log2 transformed and used for further analysis.

#### Environmental responses

The transcriptional response to heat stress was determined by explaining the gene expression over the treatment with a linear model,

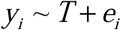

where y is the log2-normalized intensity as measured by microarray of spot i (i = 1, 2, …, 45220), and T is the treatment (either control, heat stress, or recovery from heat stress). This analysis ignored genotype.

The significances were corrected for multiple testing by applying the Benjamini Yekutieli method in p.adjust (R, version 3.3.1 Windows x64) at FDR = 0.05 [27]. Thresholds of −log10(p) ≥ 2.87 for the control versus heat-stress treatment, −log10(p) ≥ 3.09 for the control versus recovery treatment, and −log10(p) ≥ 3.02 for the heat-stress versus recovery treatment were determined.

#### Developmental variation

Due to the setup of our experiment, potential variation in development could exist among the RILs and the treatments. The recovery animals were sampled two hours later than the control and heat-stress animals, furthermore, heat stress slows the developmental rate [19]. We estimated the relative age by using a set of ∼100 genes that show a strong, positive, linear response during development [9]. By setting the average age of the control RILs to 48 hours we could estimate and compare the RILs in all treatments (**Additional file 8**).

#### Principal component analysis

A principal component analysis was conducted on the gene-expression data of the RILs over the three treatments. For this purpose, the data was transformed to a log2 ratio with the mean, using

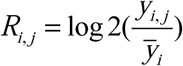

where R is the log2 relative expression of spot i (i = 1, 2, …, 45220) in strain j (RIL) over all three conditions (n = 48 per condition), and *y* is the intensity (not the log2-transformed intensity) of spot i in strain j.

The transformed data was used in a principal component analysis, where the first six axes were further examined.

### Expression quantitative trait locus analysis

#### eQTL mapping and threshold determination

The eQTL mapping was done in “R” (version 3.3.1 Windows x64). The gene-expression data was fitted to the linear model,

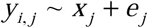

where y is the log2-normalized intensity as measured by microarray of spot i (i = 1, 2, …, 45220) of RIL j. This is explained over the genotype (either CB4856 or N2) on marker location x (x = 1, 2, …, 729) of RIL j.

The genome-wide significance threshold was determined via permutation, where the log2-normalized intensities were randomly distributed per gene over the genotypes. The randomized data was tested using the same model as for the eQTL mapping. This was repeated for ten randomized datasets. A false discovery rate was used to determine the threshold (as recommended for multiple testing under dependency) [27],

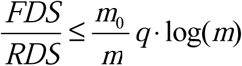

where FDS is the outcome of the permutations and RDS is the outcome of the eQTL mapping at a specific significance level. The value of m_0_, the number of true null hypotheses tested, was 45220-RDS, and for the value of m, the number of hypotheses tested, the number of spots (45220) was taken. The q-value was set at 0.05. This yielded a threshold of –log10(p) > 3.9 for the control, –log10(p) > 3.5 for the heat stress, and–log10(p) > 3.9 for the recovery treatment. For the analyses we used the most conservative thresholds measured, −log10(p) > 3.9, for all the sets.

#### Statistical power calculations

In order to determine the statistical power at the set FDR threshold, QTL were simulated using the genetic map of the strains used per condition (n = 48 per condition). For each marker location, ten QTL were simulated that explained 20-80% of the variation (in increments of 5%). Random variation was introduced based on a normal distribution with sigma = 1 and mu = 0 and a peak of the corresponding size (e.g. a peak size of 1 corresponds to 20% explained variation) was simulated in this random variation. From the simulation, the number of correctly detected QTL, the number of false positives and the number of undetected QTL were counted. This was based on the thresholds determined in the permutations, –log10(p) > 3.9. Furthermore, the precision of the effect-size estimation and the precision of the QTL location (based on a –log10(p) drop of 1.5 compared to the peak) were determined. A table summarizing the results can be found in **Additional file 9**.

#### eQTL analysis

The distinction between *cis-* and *trans-*eQTL was made on the distance between the physical location of the gene and the location of the eQTL-peak. For *cis-*eQTL the gene lies within 1 Mb of the peak or within the confidence interval of the eQTL. The confidence interval was based on a –log10(p) drop of 1.5 compared to the peak.

The amount of variation explained per microarray spot with an eQTL was calculated by ANOVA, by analysis of the gene expression explained over the peak-marker. For spots with multiple peaks, this analysis was conducted per peak, not using a full model, since a single-marker model was used in the analysis.

In order to identify *trans*-bands (an enrichment of *trans*-eQTL), a Poisson distribution of the mapped *trans*-eQTL was assumed (as in [28]). Therefore the number of *trans*-eQTL per 0.5 Mb bin were counted. Since *trans*-eQTL peaks were mapped to 107, 106, and 103 bins (respectively in control, heat stress, and recovery), it was expected that 9.16, 20.64, and 9.01 spots with a *trans*-eQTL were to be found at each of these markers. Based on a Poisson distribution, it was calculated how many *trans*-eQTL needed to be found to represent an overrepresentation. For example, for p < 0.001 there should be 20, 36, or 20 spots with a *trans*-eQTL at a specific marker (respectively in control, heat stress, and recovery).

To test for polymorphisms in genes with eQTL, we used the data from the CB4856 reference genome [22]. The genes with eQTL were matched to the polymorphisms. The frequencies of polymorphisms in each of the groups (genes with *cis*-eQTL, genes with *trans*-eQTL, and genes without eQTL) were counted and compared versus each other by a chi-squared test in “R” (version 3.3.1, x64).

#### Detection of eQTL across treatments

Two criteria were used to detect the occurrence of eQTL over multiple treatments.

In the first criterion, it was tested whether or not an eQTL was mapped in treatment one versus treatment two, by simply comparing the tables listing the eQTL. This allowed for comparison of the actual mapped peaks and for comparison of eQTL effects of *trans*-eQTL regulated from different loci. In order to estimate the false-discovery rate associated with this comparison, the same analysis was applied to ten permutated datasets per condition, using the –log10(p) > 3.9 for eQTL discovery.

The second criterion compared the occurrence of eQTL at the exact same marker location. In this comparison, the eQTL mapped in one treatment were taken as lead for the occurrence of the same eQTL in the other two treatments. This comparison allowed for direct comparison of the eQTL effect at the locus. Based on observations on the effect distribution, this approach was used to estimate the number of *trans*-eQTL not detected due to statistical power or not detected due to absence of the eQTL in a treatment (see also text in **Additional file 15**).

#### Functional enrichment analysis

Gene group enrichment analysis was done using a hypergeometric test and several databases with annotations. The databases used were: the WS220 gene class annotations, the WS256 GO-annotation, anatomy terms, phenotypes, RNAi phenotypes, developmental stage expression, and disease related genes (www.wormbase.org) [29]; the MODENCODE release 32 transcription factor binding sites (www.modencode.org) [30, 31], which were mapped to transcription start sites (according to [32]); and the KEGG pathway release 65.0 (Kyoto Encyclopedia of Genes and Genomes, www.genome.jp/kegg/) [33].

Enrichments were selected based on the following criteria: size of the category n > 3, size of the overlap n > 2. The overlap was tested using a hypergeometric test, of which the p-values were corrected for multiple testing using Bonferroni correction (as provided by p.adjust in R, 3.3.1, x64). Enrichments were calculated based on gene names, not on spots.

## Results

### Transcriptional response over the course of heat stress

To better understand the transcriptional response to heat stress, we obtained the transcriptomes of 48 recombinant inbred lines (RILs) at the L4 stage in each of three treatments: control, heat stress, and recovery from heat stress (Figure 1A). The effects of the treatments on gene-expression levels were analyzed using a linear model for pairwise comparisons between each of the conditions (see volcano plots in **Additional file 4** and a list of affected spots in **Additional file 5**). In this way, we identified 7720 differentially expressed genes over the course of the three treatments (FDR = 0.05; Figure 1B). We found that both control and heat stress had many unique differentially expressed genes: 2321 genes were only differently expressed in the comparisons of the control treatment to the other two treatments and 3305 genes were only differently expressed in the comparisons of the heat-stress treatment with the other two treatments. In the comparisons with the recovery treatment, only 942 genes were found unique for that treatment. Furthermore, many differentially expressed genes were shared in the comparison of the recovery treatment versus the control and heat-stress treatment (2251 genes). Again, the control and recovery (1152) and heat stress and recovery (1189) shared fewer genes. There were 1255 genes that were differentially expressed between all three conditions and were therefore highly treatment dependent. These results indicate that the control and heat-stress treatments are strongly contrasting in gene expression, whereas the recovery treatment shares characteristics with both other treatments.

**Figure 1:**
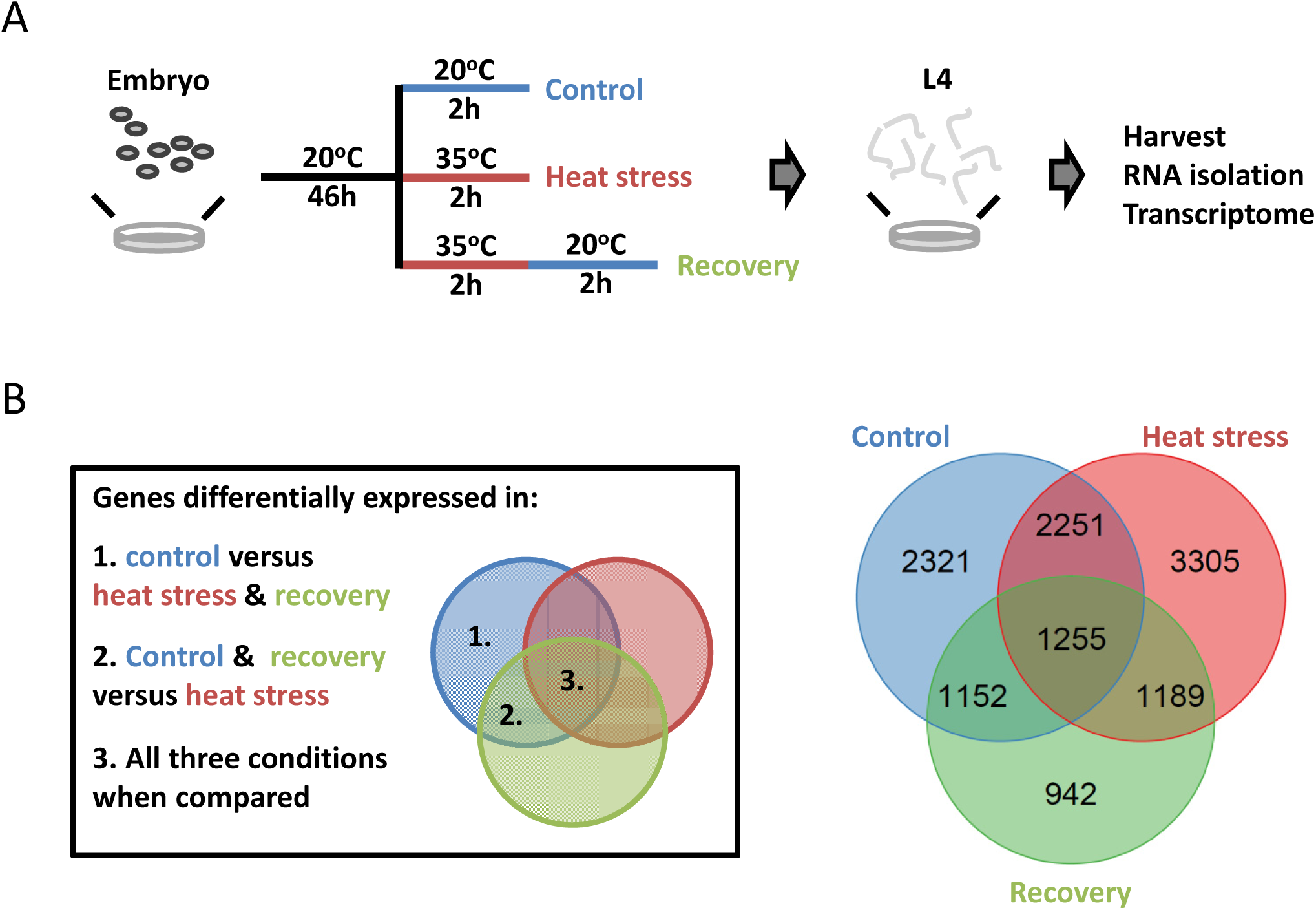
Effect of treatment on gene expression. (**A**) The RIL populations were exposed to three treatments: control (48 h at 20°C), heat stress (46 h at 20°C and 2 h at 35°C), and recovery (as heat stress, with an additional 2 h at 20°C). At the end of these treatments, the nematodes were in the L4 stage, and were harvested. Thereafter RNA was isolated and the transcriptome was measured by microarray. (**B**) The outcome of the treatment analysis. On the left a legend is included to clarify which contrasts are compared. On the right, the overlap in differentially expressed genes is shown per treatment comparison. For example, 942 genes are uniquely differentially expressed in the recovery treatment, these genes are differently expressed between recovery and heat stress and between recovery and control, but not between heat stress and control.

To follow up on this interpretation, a principal component analysis (PCA) was conducted on the gene-expression data, transformed to the log2 ratio with the mean. The first axis (20.0% of the variation) captures variation mostly related to the heat-shock treatment, which is expected as this treatment specifically affect the largest number of genes (**Additional file 6**). The second axis (11.4% of the variation) captures variation mostly related to the control treatment, which also fits the analysis with the linear model as the control treatment was the second most distinct treatment. Together, these two axes also place the recovery treatment in between the control and heat-stress treatment, showing the contrast with the other two treatments was lower and possibly indicating transcript levels in the recovery treatment are returning to normal. The third principal component (10.1% of the variation) captures variation that sets the heat-stress treatment completely apart from the other two treatments, which is as expected since the heat-stress treatment has the most unique differentially expressed genes.

In order to gain further insight into the functional differences between the treatments, an enrichment analysis was conducted on genes belonging to the different overlap groups as shown in Figure 1 (For example genes differentially expressed in only the control treatment, see the list in **Additional file 7**). Each of the groups was enriched for many processes, showing that the treatments had a profound impact on gene expression. Interestingly, we found that genes specific for each of the three treatments were enriched for genes expressed in the intestine. Furthermore, the control and heat-stress treatments were strongly enriched for genes expressed in the germline. These enrichments indicated that the expression of genes involved in metabolism and reproduction (or development of the reproductive organs) were strongly altered during the heat-stress response. As this may be caused by a developmental difference in the sampled populations, we estimated the developmental age using a transcriptional ruler (see **Materials and methods** for details) [9]. It was found that the control population was transcriptionally slightly younger than the heat shock (estimated ∼1 h older) and the recovery population (estimated 1.3 h older; **Additional file 8**).

### Gene expression linked to genetic variation

Linkage mapping was performed using 48 RILs for each of the three treatments. Statistical power analysis showed that this population has the power to detect 80% of the eQTL that explain at least 35% of the variation (see Materials and methods and the table in **Additional file 9**). Identified eQTL (FDR ≤ 0.05; Figure 2; a table with all eQTL is given in **Additional file 10**) were compared between the treatments (Table 1). Most genes with an eQTL were found in the heat-stress treatment (2626), whereas the control (1797) and recovery (1880) had similar numbers. This increase in genes with eQTL was primarily caused by the larger number of genes with *trans*-eQTL in the heat-stress treatment (1560; ∼57% of total eQTL) compared to the control (751; ∼40% of total) and recovery (739; ∼38% of total). The number of *cis*-eQTL was almost identical among conditions (Table 1).

**Figure 2:**
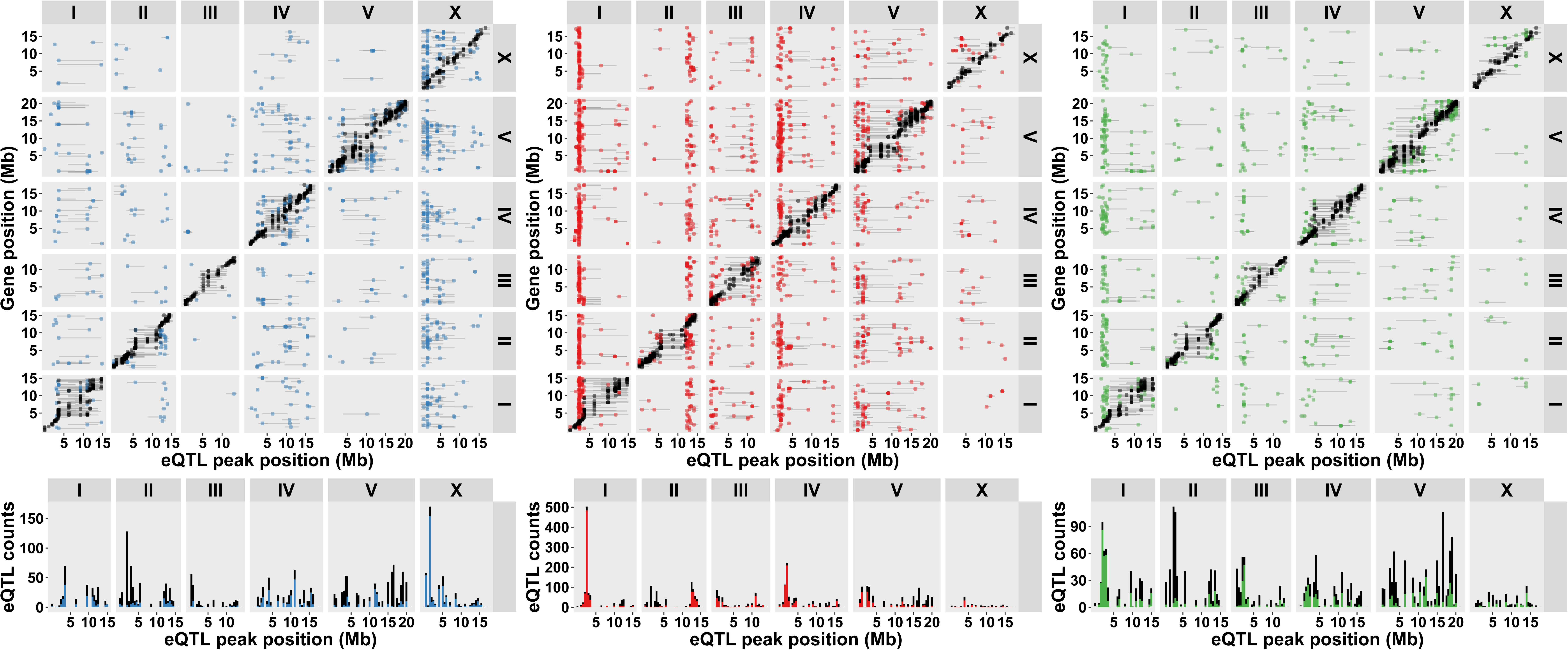
**Identified eQTL** in control (left), heat stress (middle), and recovery (right) treatments, with a threshold of −log10(p) > 3.9 (FDR ≤ 0.05) in each treatment. The eQTL peak position is shown on the x-axis and gene position is shown on the y-axis. The *cis*-eQTLs (within 1Mb of the gene) are shown in black and the *trans*-eQTLs in blue (control), red (heat stress), or green (recovery). The horizontal bars indicate the confidence interval of the eQTL. The chromosomes are indicated on the top and right of the plot. The histogram under the plot shows the eQTL density per 0.5 Mb bin.

**Table 1:** Number of genes with an eQTL

| **Table 1: Number of genes with an eQTL** |  |  |  |
| --- | --- | --- | --- |
|  | **Control** | **Heat stress** | **Recovery** |
| *cis*-eQTL<sup>1</sup> | 1138 | 1186 | 1215 |
| N2 higher | 809 (71.1%) | 802 (67.6%) | 883 (72.7%) |
| CB4856 higher | 329 (29.9%) | 384 (32.4%) | 332 (27.3%) |
| *trans*-eQTL | 751 | 1560 | 739 |
| N2 higher | 300 (39.9%) | 583 (37.4%) | 393 (53.2%) |
| CB4856 higher | 451 (60.1%) | 977 (62.6%) | 346 (46.8%) |
| Total<sup>2</sup> | 1797 | 2626 | 1880 |
| <sup>1</sup>: *cis*-eQTL were called if the QTL peak lies within 1 Mb of the affected gene, or if the affected gene lies within the 1.5 LOD-drop confidence interval.<br><sup>2</sup>: The differences between the total and the summation of the individual *cis*- and *trans*-eQTL is due to genes with both a *cis*- and *trans*-eQTL. |  |  |  |

The *cis*-eQTL showed a bias for higher expression if the regulatory locus had the N2 allele (on average 70% of the genes with a *cis*-eQTL). This bias was absent in the *trans*-eQTL where, on average, 43% of the genes with a *trans*-eQTL was more highly expressed if the locus had the N2 allele (illustrated by the figure in **Additional file 11**). This discrepancy was to be expected, since the microarray platform used to measure the transcripts was designed for the N2 genotype. Therefore, part of this variation is likely due to mis-hybridization. However, previous studies have shown that this does not explain all the variation (see [28]). In congruency, we found genes with a *cis*-eQTL to be more polymorphic than genes with a *trans*-eQTL and genes without an eQTL (summarized in **Additional file 12**). For example, genes with a *cis*-eQTL were more likely to be fully deleted in CB4856 (9.5% of the genes, compared to 0.9% in genes without an eQTL or 1.1% in genes with a *trans*-eQTL; Chi-squared test, P < 1*10^−34^). Furthermore, supporting that not all *cis*-eQTL stem from mis-hybridization, polymorphisms in the flanking 3’ and 5’ regions were about two times more likely to occur near *cis*-eQTL compared to genes without a *cis*-eQTL (Chi-squared test, P < 1*10^−6^). When these enrichments for 3’ and 5’ polymorphisms were compared between *cis*-eQTL with an N2 or CB4856 higher effect, we found no significant difference.

The *trans*-eQTL were mainly found in 19 treatment-specific *trans*-bands, loci that regulate the abundance of many transcripts. The 19 *trans*-bands were identified by analysis of the occurrence of *trans*-eQTL across the genome (Poisson distribution, P < 0.001; listed in **Additional file 13**). The seven *trans-*bands detected in the heat-stress treatment affected most genes (1141; ∼73% of all *trans*-eQTL). In the control treatment, five *trans*-bands were found (325 genes; ∼43% of all *trans*-eQTL) and in recovery treatment seven *trans*-bands were identified (343 genes; ∼46% of all *trans*-eQTL). Six out of the 19 *trans*-bands individually affected >100 genes, one located at chromosome X: 0.5-2.0 Mb in control treatment; four in heat-stress treatment at chromosome I: 2.0-3.5 Mb, II: 12.0-13.5 Mb, IV: 1.0-2.5 Mb, and V:1.0-3.0 Mb; and one in recovery treatment at chromosome I: 1.5-3.0 Mb. Importantly, the distribution across the genome of *trans*-bands and eQTL is treatment-specific. To further investigate this, we determined the overlap in mapped eQTL over the course of the heat-stress response.

### In contrast to *cis*-eQTL, trans-eQTL were environment-specific

Comparing the genes with a *cis*-eQTL among the three treatments, we found 1086 out of 1789 unique genes (∼61%) with a *cis*-eQTL in more than one treatment and 664 (∼37%) in all three treatments (Figure 3A). Because *cis*-eQTL can be caused by mis-hybridizations, we also calculated the overlap for *cis*-eQTL with an N2 higher effect. In that selection, 303 out of 615 unique genes (∼49%) had a *cis*-eQTL in more than one treatment and 157 (∼26%) in all three treatments. For the whole set of genes with a *trans*-eQTL the overlap was much smaller (Figure 3B); 360 out of 2610 genes (∼14%) were found in more than one treatment and only 80 genes (∼3%) in all three treatments. By definition, the locus at which *cis*-eQTL were detected was the same between treatments. Furthermore, the *cis*-eQTL effect sizes and directions were highly comparable (Pearson correlation coefficients between 0.94-0.96; shown in a figure in **Additional file 14**). The *cis*-eQTL that were detected in only one treatment were probably missed due to the small amount of variation explained by these eQTL (see the text in **Additional file 15**). Interestingly, there were only three genes with *cis*-eQTL showing genotypic plasticity: C52E2.4, C54D10.9, and *nhr-226*. This meant that we observed allelic variation acting in opposite directions between environments (Figure 3C).

**Figure 3:**
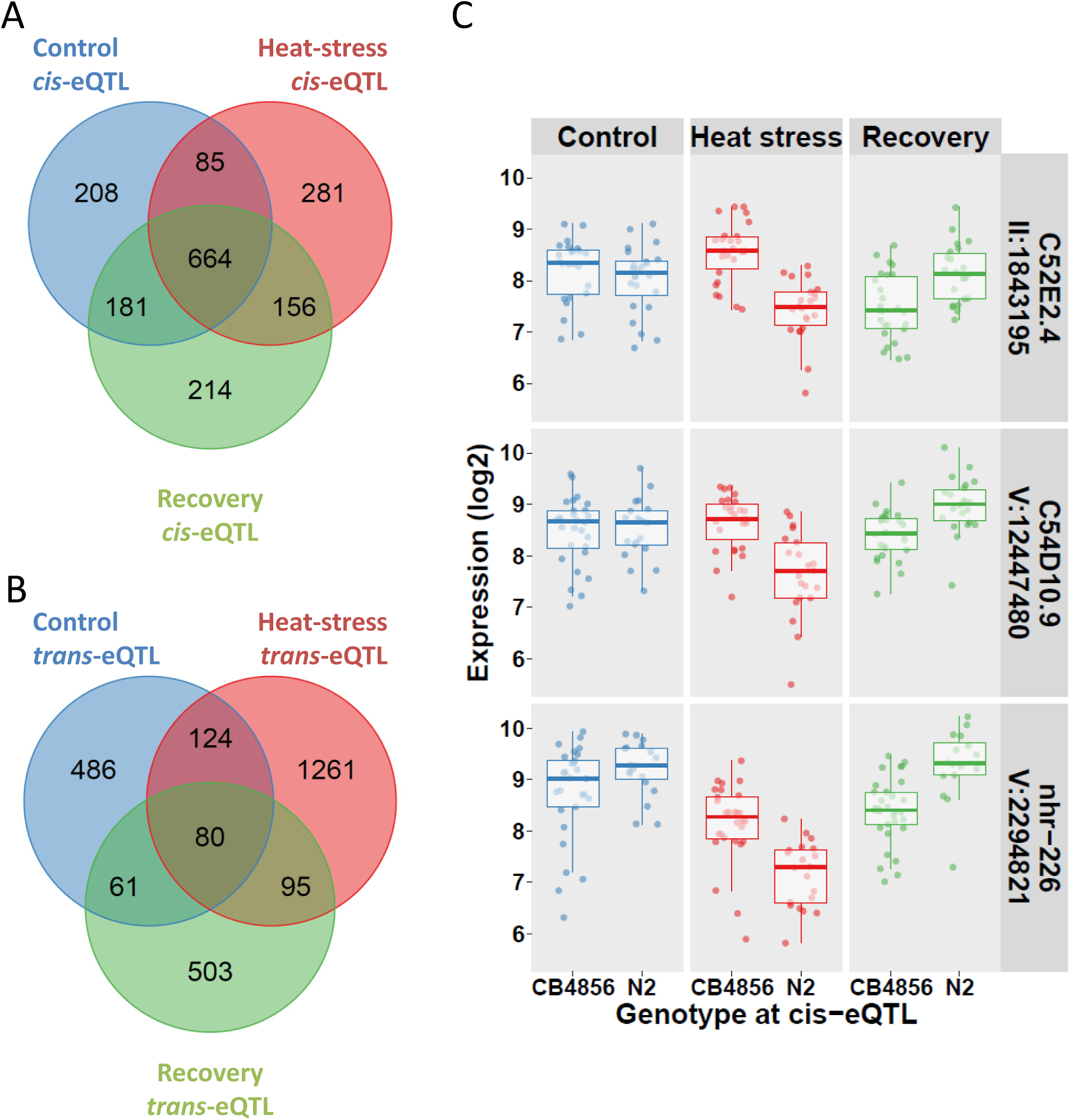
Overlap in *cis*- and *trans*- eQTL between conditions. (**A**) The overlap in genes with a *cis*-eQTL between conditions. QTL were selected based on FDR ≤ 0.05 per condition, a QTL was scored as overlapping if the same gene had a *cis*-eQTL in another condition. (**B**) The overlap in genes with a *trans*-eQTL between conditions. QTL were selected based on FDR ≤ 0.05 per condition, a QTL was scored as overlapping if the same gene had a *trans*-eQTL in another condition. The location was not yet considered in this analysis (see **Supplementary figure 7A**). (**C**) The three genes with a *cis*-eQTL displaying genotypic plasticity, where the direction of the effect switches between environments. For all three genes: C52E2.4, C54D10.9, and *nhr-226* the plastic response is apparent between the heat stress and recovery treatment.

The effect sizes and direction of genes with a *trans*-eQTL found in more than one treatment were very similar (Pearson correlation coefficients between 0.90-0.93; figure in **Additional file 14**). However, only a few genes with a *trans*-eQTL were found over multiple treatments, far lower than the overlap in *cis*-eQTL between conditions (Figure 3A and B). Since *trans*-eQTL are not by definition regulated from the same location, the overlap between treatments declines even further if location is taken into account. Taking the *trans*-eQTL from the three treatments together, only ∼38% of the multi-treatment *trans*-eQTL were located at a different locus from one treatment to another (∼28% if only loci at different chromosomes were counted), see **Additional file 16** (figure A). Interestingly, although the regulatory locus was located elsewhere, the genotypic effect of the *trans*-eQTL was almost identical (**Additional file 16,** figure B), yet most genes with *trans*-eQTL only displayed an eQTL in one treatment (text in **Additional file 15**). The likely reason that the majority of *trans*-eQTL was not detected across treatments is that most *trans*-eQTL are environment-specific and therefore highly cryptic.

### *Trans*-eQTL display two types of cryptic variation

For treatment-specific, cryptic *trans*-eQTL, we found that a regulator is active when the eQTL is detected and it is not active when no eQTL is detected. With on and off switching regulators between treatments, genes can have dynamic *trans*-eQTL, which appear as a treatment-specific switch of regulatory loci (Figure 3 – 5). Since the majority of genes with a *trans*-eQTL have one unique *trans*-eQTL in only one treatment (Figure 3B), a switch in regulatory loci seems to occur less frequently compared to the on/off switch.

As the majority of the *trans*-eQTL was treatment-specific, we investigated whether detection in only one treatment was a result of the statistical power of our study or if it was a biological phenomenon. As *trans*-eQTL explain 34.2% of variation on average, compared to 52.1% of variation for *cis*-eQTL (text in **Additional file 15**), it is possible that the detection of *trans*-eQTL was more affected by lack of statistical power. By simulations, we estimated that this only affected 20.6% of the *trans*-eQTL that were not detected in multiple treatments (for the detailed analysis, see the text in **Additional file 15**), which argues for the cryptic nature of *trans*-eQTL. Another line of evidence for this is the low overlap in affected genes in co-locating *trans*-bands across treatments (13/19 *trans*-bands co-locate). We only found significant overlap in three pairs of *trans*-bands, where 6.5-11.1% of the affected genes overlap (text in **Additional file 15;** hypergeometric test p < 1*10^−4^). For example, one of these, a major *trans*-band at chromosome IV:1-2.5 Mb in the heat-stress treatment, affected 244 genes of which 22 overlapped with the 31 genes in the recovery *trans*-band on chromosome IV:1-2 Mb. Together, these results show that the majority (67.3%) of *trans*-eQTL indeed are cryptic. For example, the transcript levels of *flp-22*, *pqm-1*, and *sod-5* were affected by treatment (Figure 4A) and showed a *trans*-eQTL in only one treatment (Figure 4B).

**Figure 4:**
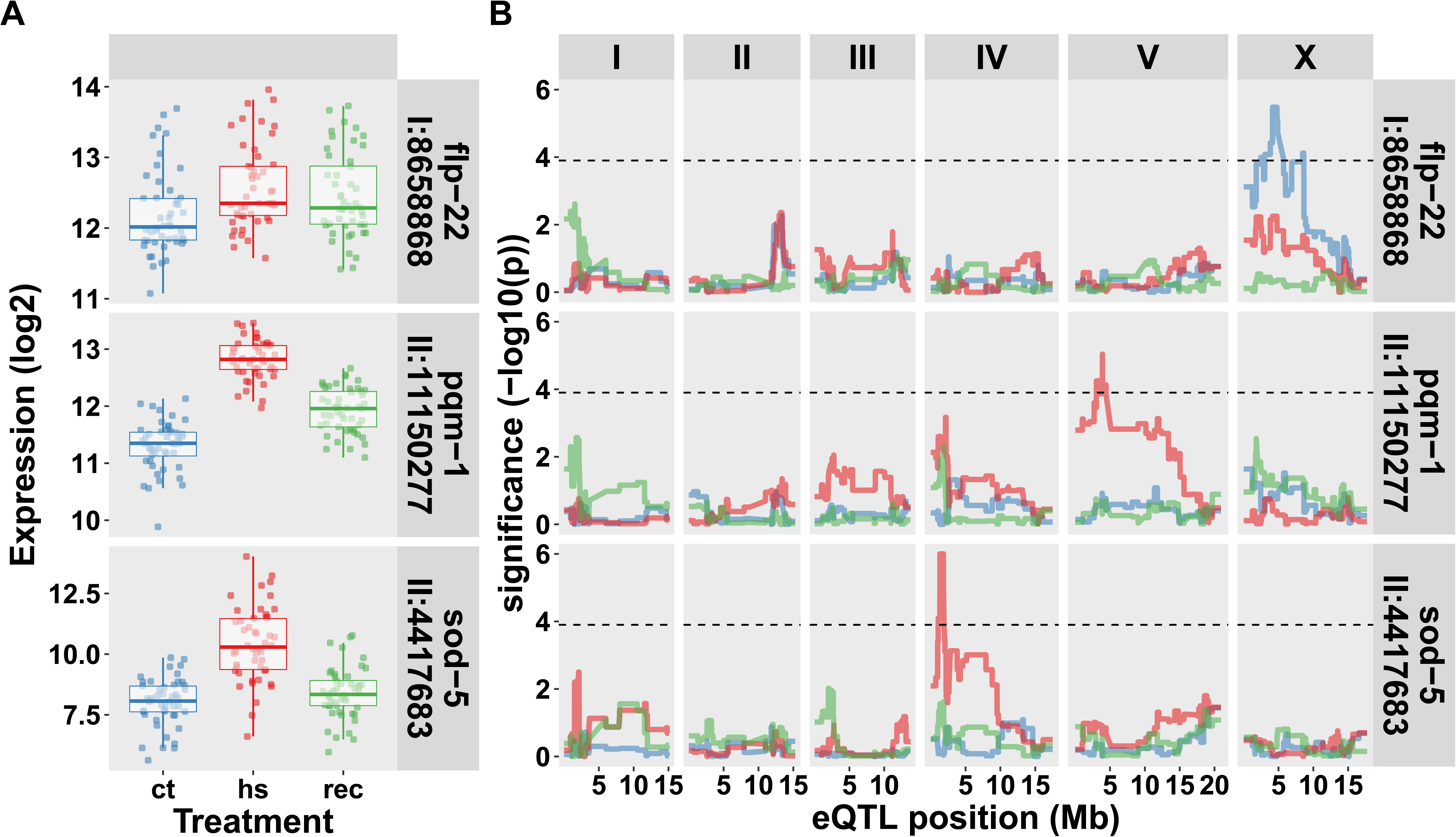
Genes with treatment specific *trans-*eQTL. (**A**) The expression patterns over the three treatments are shown for *flp-22*, *pqm-1*, and *sod-5*, the location mentioned is the location of the gene. The x-axis is organized per treatment (ct, control; hs, heat stress; rec, recovery). On the y-axis the log2 normalized expression is shown (**B**) The eQTL patterns for the same three genes. The x-axis shows the position along the chromosomes and the y-axis the significance of the association. The horizontal dashed line indicates the FDR ≤ 0.05 (−log10(p) > 3.9). Colors indicate the three treatments.

As mentioned before, only a minority of the genes with a *trans*-eQTL have different eQTL across treatments. Of those genes with different *trans*-eQTL over treatments, the genotypic effects of the eQTL were similar across treatments, even if the loci were different (**Additional file 16,** figure B). Only between chromosome IV and V were eQTL with changes in effect directions observed. One of the genes displaying this pattern was *gei-7* (also known as *icl-1*). This gene was represented by three different micro-array probes, all showing the same pattern: a primary eQTL on chromosome V at ∼12.0Mb in all three conditions and a secondary eQTL in the heat-stress treatment at chromosome IV at ∼1.5Mb (Figure 5).

**Figure 5:**
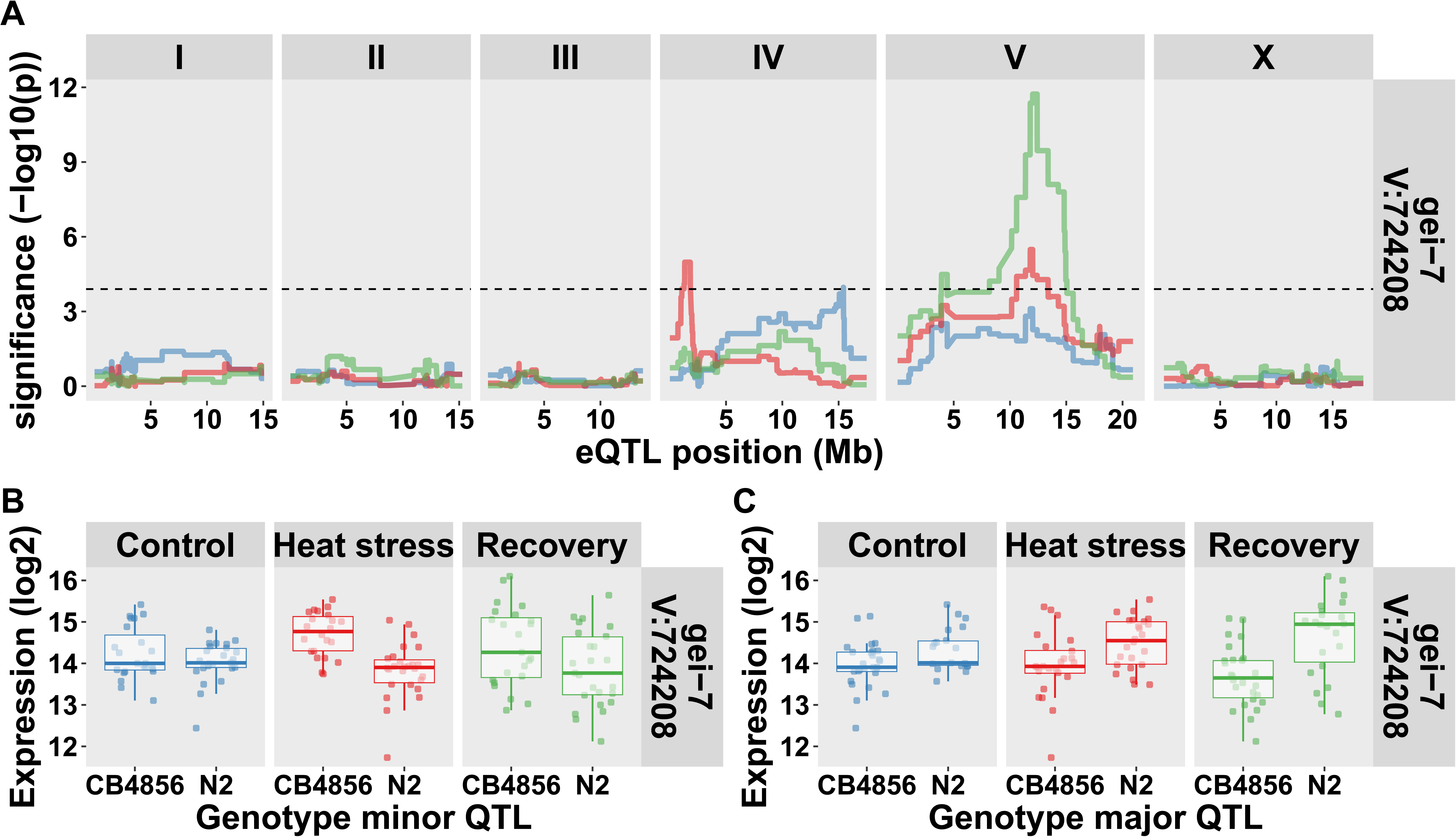
The *trans*-eQTL of *gei-7*. (**A**) The eQTL patterns for *gei-7*, the x-axis shows the position along the chromosomes and the y-axis the significance of the association. The horizontal dashed line indicates the FDR ≤ 0.05 (−log10(p) > 3.9). Colors indicate the three treatments (control, blue; heat stress, red; recovery, green). (**B**) The genotype effects split out at the minor heat stress QTL (chromosome IV). (**C**) The genotype effects split out at the major QTL (chromosome V).

### Functional enrichment of eQTL

To find which biological processes were affected by genetic variation on a gene-expression level we looked for enrichment in gene classes, phenotypes, KEGG pathways, GO terms, and anatomy terms (see the list in **Additional file 17**). For *cis*-eQTL enriched categories were similar in all three treatments, as expected by the consistent nature of *cis*-eQTL. For *cis*-eQTL, we found enrichments for the gene classes *bath*, *math*, *btb*, and *fbxa*, which were previously found to be highly polymorphic between CB4856 and N2 [22]. Moreover, we found enrichment for genes involved in the innate immune response and protein homo-oligomerization. It should be noted that these enrichments are likely due to hybridization differences for the polymorphic genes (as *cis*-eQTL with a positive N2 effect are enriched for exactly these categories, **Additional file 17**).

The *trans*-eQTL were also enriched for genes functioning in the innate immune response, especially for genes where the N2 allele leads to higher expression. Furthermore, genes expressed in the intestine were enriched in the *trans*-eQTL found in control and heat-stress conditions. Contrasting to the genes with *cis*-eQTL, the genes with *trans*-eQTL were enriched for many different transcription factor binding sites, indicating active regulation of *trans*-eQTL.

Consistent *trans*-eQTL were found in all three treatments for the enriched NSPC (nematode specific peptide family, group C) gene class. This was remarkable as only a very small part of the *trans*-eQTL were shared over the three treatments. For the heat stress and recovery *trans*-eQTL, genes expressed in the dopaminergic neuron were enriched, with the strongest enrichment in the heat-stress treatment. These genes were also enriched in the control treatment, however, in a group of *trans*-eQTL mapping to a different trans-band. In the heat stress and recovery treatment the dopaminergic neuron-specific genes showed *trans*-eQTL at the IV:1-2MB locus, whereas in the control they showed *trans*-eQTL at the X:4-6 locus (Figure 6).

**Figure 6:**
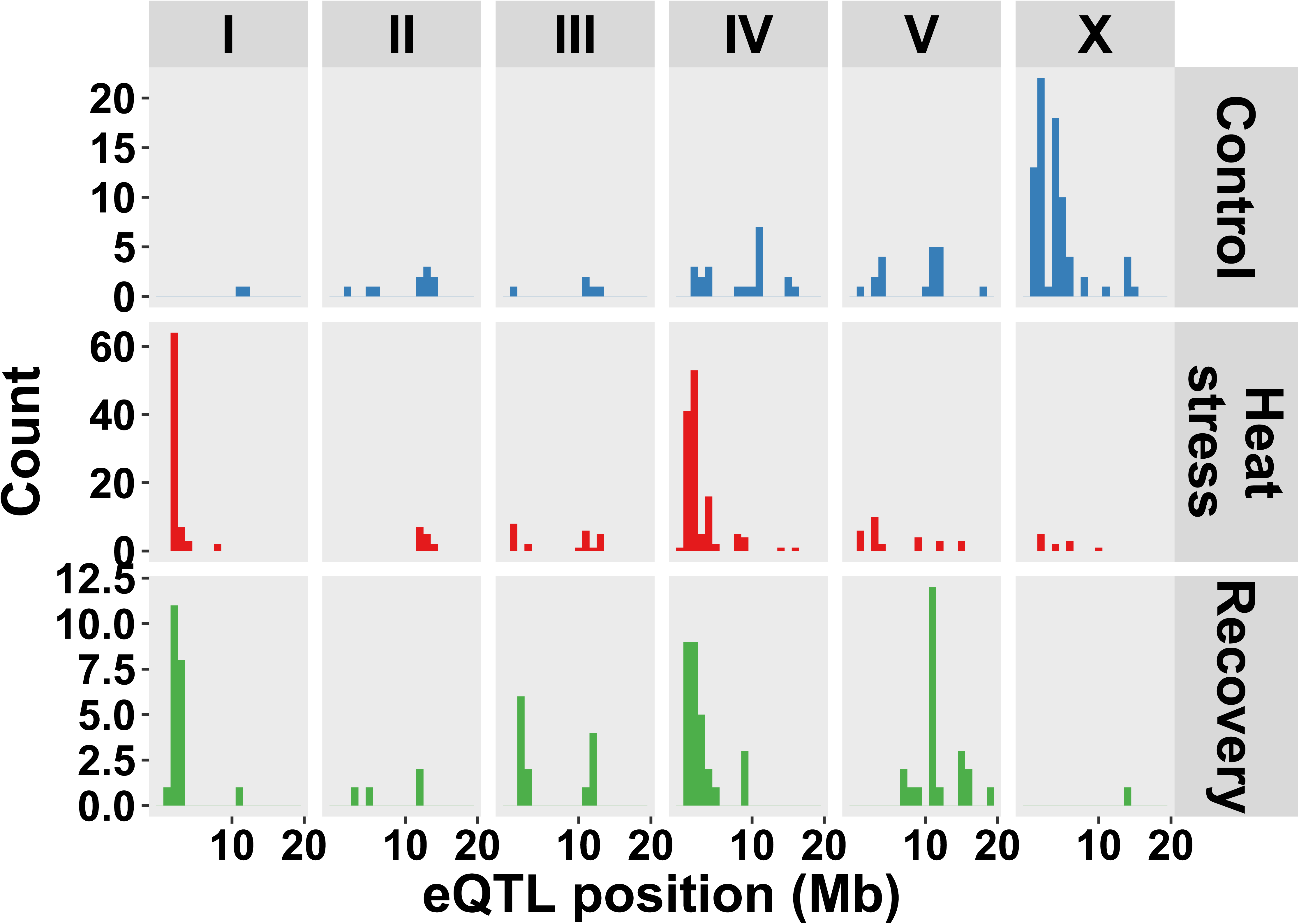
Genes belonging to the dopaminergic neuron anatomy term with a *trans*-eQTL. The *trans*-eQTL position of the genes is shown (n = 133 in control, n = 281 in heat stress, and n = 91 in recovery).

## Discussion

### Transcriptional response over the course of a heat stress

Here we present a comprehensive study of the effects of an induced heat-stress treatment on the genetic architecture of gene expression. The obtained transcriptomes were analyzed in the context of the treatments and in the context of the genetic variation present in the strains used in the experiments.

We found that many of the genes that are affected over the heat-stress course are associated with expression in the germline and intestine. These findings are partially in line with findings from an investigation on the heat-shock regulatory network using a genome-wide RNAi screen in *C. elegans* [5]. Their heat-stress conditions were 31.5°C for two hours followed by 24 hours of recovery at 20°C. The authors found that genes associated with the proteasome induced heat-stress-specific gene expression only in the intestine and spermatheca, which corroborates with our results. Differences with Guisbert *et al.* (2013), could be explained by the differences in larval stage used (L4 vs L2) as well as differences in temperature and duration of the heat stress (and recovery). Another study exposed *C. elegans* at the L4 stage to a heat stress of 30 min at 33°C, and measured transcriptome differences using RNA-seq [34]. In support of our study, they also detected genes associated with metabolism and reproduction, whereas, in contrast to our findings, they found a strong link with cuticle specific genes. This contrast could be due to the different experimental conditions of Bunquell *et al.* (2016), as they used RNAi treatments (empty vector or against *hsf-1*), a different heat shock duration, method of synchronization (additional L1 arrest by [34]), and rearing temperature before heat shock (23°C versus 20°C in this study).

We hypothesize that ultimately these discrepancies are likely to result from developmental differences. The transcriptional program in *C. elegans* differs strongly during development [8, 9]. Therefore, application of a heat shock on L2 larvae has a very different developmental (and therefore transcriptional) starting point compared to a heat shock applied on L4 larvae. Furthermore, within the L4 stage there is a strong difference in gene expression in early-stage L4 and late-stage L4. We estimated that the heat-stress and recovery RILs were one hour older than the control RILs suggesting that the former had an accelerated development as these populations were physically the same age as the control population. The main processes that affect transcription in the L4 stage are reproduction and development [9]. These are exactly the processes that are halted upon induction of a heat shock [5, 34]. Therefore, it is likely that the state in which these processes are strongly affects the possible routes for down regulation. It would therefore be very interesting to study the effect of heat shock in relation to the developmental dynamics.

If the effect of a heat shock is indeed dependent on developmental status, then the transcriptomes of the RILs presented in this study could be indicative of the phenotypic outcome (*e.g.* heat-stress survival). The reason being the developmental gradient generated by RILs (as shown by [15] within a single experiment, or explicitly by [35] between different stages), which mainly affects *trans*-eQTL. Alternatively, our experiment could also be analysed by including the variation among the estimated age of the RILs in the mapping model together with treatment effects. A thorough interpretation of these models requires a better understanding of the transcriptional dynamics over the heat-shock and recovery response, a goal we are currently actively pursuing [36].

### In contrast to *trans*-eQTL, *cis*-eQTL are directly linked to polymorphisms

The *cis*-eQTL over the three treatments, which strongly overlapped, are highly enriched for polymorphic genes. This has been reported before in *C. elegans*, but also in *A. thaliana*, *Mus musculus*, and for human *cis*-eQTL [12, 28, 37–41]. This can result in the detection of transcriptional variation that is actually caused by hybridization differences [28, 40]. Analysis of the bias in *cis*-eQTL with higher expression in N2 (the strain for which the microarray was developed) versus CB4856 indeed shows that a proportion of the *cis*-eQTL is likely to stem from hybridization differences. Also apparent from the gene-enrichment analysis, *cis*-eQTL were overrepresented for polymorphic gene classes such as *bath*, *math*, *btb*, and *fbxa*, which are also divergent among other wild strains [42, 43]. Other experimental methods could limit such ‘false positives’, for example, RNA sequencing is expected not to suffer from such biases [44].

Interestingly, genes with a *cis*-eQTL were also strongly enriched for polymorphisms in regulatory regions. For these *cis*-eQTL, it could be true that the expression is affected by transcription factor (TF) binding sites [45], yet we did not detect any enrichment for such sites as mapped by ModEncode [30, 31]. An explanation for this is that genes with a *cis*-eQTL are regulated by different TFs, therefore the affected TF binding site is different per gene with a *cis*-eQTL making an overrepresentation among all *cis*-eQTL unlikely.

### CGV of transcriptional architecture is determined by *trans*-eQTL

Although previous studies in *C. elegans* have focused on continuous (thermal or developmental) treatments or gradual change over time [12, 15, 35], only few genetical genomics experiments have measured the effect of acute perturbation, where an organism is suddenly exposed to a different environment [39, 46]. This acutely affected the transcriptional architecture of gene expression, which consists of *cis*- and *trans*-acting eQTL. We found that *cis*-eQTL were robust across all three treatments, including heat stress, which was found before in studies on *C. elegans* and other species where eQTL patterns have been studied in different conditions. For example, Li *et al.* (2006) found that more than 50% of all *trans*-eQTL were affected by temperature compared to *cis-*eQTL [12], and Smith and Kruglyak (2008) also reported that *cis*-eQTLs were hardly affected by external conditions as compared to *trans*-eQTL [47]. Also, in other species, like *Arabidopsis thaliana*, it was reported that *cis*-eQTL were robust to light perturbation, whereas *trans*-eQTL were more affected by different light regimes [39]. In humans, *cis*-eQTL showed a very high correlation between tissues compared to *trans*-eQTL, and are replicated across populations in lymphoblastoid cell lines [48, 49].

By inducing a strong environmental perturbation in the form of a heat shock, a high amount of different *trans*-eQTL were detected across the control, heat-stress, and recovery treatments. One of the things we noted is the increase in the number of *trans*-eQTL and *trans*-bands in the heat-stress treatment compared to the other two treatments. This strong transcriptional variation could underlie – or result from - the phenotypic trait differences observed between N2 and CB4856 in temperature experiments. However, it should be noted that also developmental differences due to timing between the three treatments can contribute [9, 15]. Temperature-affected trait differences have been observed for age at maturity, fertility, body size, vulval induction, and lifespan [19, 50–53]. Furthermore, these strains also display behavioural differences in heat avoidance and thermal preference [20, 21]. Likely candidates for these trait differences could be found in loci affecting the expression of many *trans*-eQTL. For example, the left arm of chromosome IV harbours a *trans*-band affecting the expression of 244 genes and coincides with a QTL affecting lifespan after heat shock [19]. Analogously, the laboratory allele of *npr-1* affects many trait differences between N2 and CB4856 (as reviewed in [54]). Most importantly, *npr-1* affects the behaviour of the animal, which probably results in gene-expression differences, as part of the expression differences can be mimicked by starving nematodes [55]. These gene-expression differences can be picked up as a *trans*-band [28]. The latter example illustrates both a link between gene-expression and classical traits and that caution is required for inferring the direction of causality.

Why is CGV mainly affecting *trans*-eQTL? We hypothesize that this is due to the versatile nature of *trans*-eQTL. First, *trans*-eQTL are loci that are statistically associated with variation in transcript abundance from genes elsewhere on the genome. The ultimate causes for this association can be manifold, from direct interactions such as polymorphic transcription factors that affect gene expression to indirect interactions such as receptor-kinase interactions [46], receptors [46, 55, 56], or effects at the behavioural level that result in expression differences [55]. Therefore, an environment can require the organism to respond, thereby requiring specific polymorphic genes to react, ultimately leading to the expression of cryptic variation.

### The *trans*-eQTL architecture is comprised of treatment-specific genes

The *trans*-eQTL architecture is remarkably unique over the three treatments tested. We only observed 3% overlap in *trans*-eQTL in the three treatments, for which the main cause was treatment specificity of *trans*-eQTL. Surprisingly, genetic variation affects the expression of genes in only one direction; only in rare cases does allelic variation change the sign of the effect and this was only observed for *cis*-eQTL (C52E2.4, C54D10.9, and *nhr-226*). On the one hand, this is in congruency with other eQTL studies comparing different environments; *trans*-eQTL are strongly affected by different environments (for example, see [12, 39, 47]). On the other hand, it raises questions about the genetic architecture of *trans*-bands; co-localizing *trans*-bands are generally not affecting the expression of the same genes (see text in **Additional file 15**), which can imply the involvement of multiple regulators (causal genes).

However, it seems unlikely that an abundance of novel *trans*-eQTL also implies an abundance of novel causal genes. First of all, over all three treatments, the majority of *trans*-eQTL are located in *trans*-bands, which are mostly non-overlapping between treatments. This is an observation that extends to other studies and other species in which eQTL have been mapped: the majority of *trans*-eQTL are found on a few regulatory hotspots (for example, see [28, 38, 57]). Therefore, it is logical to assume that a small set of causal genes ultimately explains the majority of *trans*-eQTL. Second, the allelic effect only has one direction, which is easily aligned with the notion of a few regulators instead of many. Together, these observations can help in further dissecting loci to identify causal genes. *Trans*-band regulators might play a role in the dynamic response, which could aid in narrowing down candidate genes.

However, it should be reiterated that the route from genetic variation resulting in transcriptional effect can be manifold, which can also obscure the ultimate cause of the observed trait variation. An organism is an intricate web of interdependencies leading to the phenotype. Therefore, upon further dissection of the loci, a single eQTL might prove to be many. Furthermore, it should be noted that *trans*-eQTL explain less variation compared to *cis*-eQTL, which has been established across species [28, 58]. Therefore, it is more likely that *trans*-eQTL are not detected. However, the treatment specificity and direction of allelic effects of *trans*-eQTL across three treatments robustly show *trans*-eQTL architecture is comprised of treatment-specific genes.

## Conclusion

Here we present the contribution of CGV on eQTL across three treatments in the nematode *C. elegans*. We find that mainly *trans*-eQTL are affected by CGV, in contrast to *cis*-eQTL, which are highly similar across treatments. Furthermore, we show that most CGV results in unique genes with a *trans*-eQTL, instead of different allelic effects and/or different eQTL for the same genes. This shows the highly dynamic nature of CGV.

## Declarations

### Availability of data and materials

All strains used can be requested from the authors. The transcriptome datasets generated and the mapped eQTL profiles can be interactively accessed via (http://www.bioinformatics.nl/EleQTL). Moreover, the transcriptome datasets are also deposited at ArrayExpress (E-MTAB-5779).

### Authors’ contributions

LBS and JK conceived and designed the experiments. MGS, RPJB, RJMV, JAGR, and RB conducted the experiments. AVH, RB conducted the sequencing of the strains. LBS, MGS, and RPJB conducted transcriptome and main analyses. LBS, MGS, and JEK wrote the manuscript. RPJB, RJMV, and AC provided comments on the manuscript.

## Acknowledgements

The authors are grateful for the discussions and support from the GRAPPLE project partners: Olga Valba, Sreenivas Chavali, Benjamin Lang, Mirko Francesconi, Sergei Nechaev, Olga Vasieva, M. Madan Babu, and Ben Lehner. We are also grateful for Katharina Jovic, Lisa van Sluijs and two anonymous reviewers for feedback on the manuscript. We also want to thank Harm Nijveen for making our data available in EleQTL.

## Additional files

**Additional file 1: Strains and genotypes**. Matrix with the strain names and genotypes of the recombinant inbred lines used in this study. The genotypes are based on genome sequencing (see Materials and Methods).

**Additional file 2: A figure of the location of the 729 markers**. The marker locations are plotted across the genome. Locations are based on WS256.

**Additional file 3**: **A figure of the marker correlation analysis**. Correlations between the 729 markers in the sequenced RIL population. The markers are plotted at their physical locations across the chromosomes.

**Additional file 4: Figure of the treatment comparisons**. Volcano plots of the expression comparisons per treatment (n = 48 RILs per treatment). The horizontal line in the plots indicates the FDR = 0.05 threshold. The colored dots indicate spots that are significantly different between treatments. Blue indicates spots more highly expressed in the control treatment, red indicates spots more highly expressed in the heat-stress treatment, and green indicates spots more highly expressed in the recovery treatment. (**A**). The comparison between control and heat stress, threshold: −log10(p) ≥ 2.87. (**B**). The comparison between control and recovery, threshold: −log10(p) ≥ 3.09. (**C**). The comparison between heat stress and recovery, threshold: −log10(p) ≥ 3.02.

**Additional file 5: Table of treatment comparison results**. A table with the spots that are significantly different between treatments. The spot number, the comparison in which the spot was different, and the characteristics of the difference (significance and effect) are given. Also information about the gene represented by the spot is shown (WormBase identifier, sequence name, public name, and the location on the genome). The last column indicates to which group the spot belongs (*e.g.* specific for heat-stress treatment, or significantly different between all three treatments).

**Additional file 6: Figure of the principal component analysis**. The first six axes of the principal component analysis are shown, plotted against each other. The axis capturing most variation (PC1), captures 20% and the sixth axis captures 4.7%. The dots represent individual samples (n = 48 RILs per treatment) and are colored according to treatment: blue for control, red for heat stress, and green for recovery. The first axis separates the heat-stress treatment from the control and recovery treatment, but also captures some technical variation (19 samples that fall to the right of the plot). The second axis (11.4%) places the recovery treatment in between the other two treatments, whereas the third axis (10.1%) separates the heat stress from the control and recovery treatment.

**Additional file 7: A list of the enrichment analysis treatment responsive genes**. Enrichment analysis on the genes with transcriptional responses by heat stress. The database used for enrichment (Annotation) and the category (Group), and the number of genes on the array that are in the group (Genes_in_group) are also indicated. Furthermore, the overlap with the cluster (Overlap) and the Bonferroni-corrected significance of that overlap are shown.

**Additional file 8: Developmental ruler applied to the RIL populations.** The age of the samples was estimated using the transcriptional ruler from [9], the reported ages are relative, where the average age of the control samples were set to 48 hours. Each point represents one sample, whereof the relative age was determined by assessing the expression of about 100 genes.

**Additional file 9: Table summarizing the statistical power calculations.** Outcome of the statistical power calculations conducted for the RIL population of each treatment (n = 48 RILs per treatment). The outcomes are ordered per treatment population, and per simulated QTL peak size. All peaks were simulated in random variation generated by a standard normal distribution. The simulation reports on (i) QTL detection, *e.g.* how many of the simulated QTL were detected, how many false QTL were reported; (ii) QTL effect size estimation, mapped effect/simulated effect (reported in quantiles); (iii) QTL location estimation, |mapped location – simulated location|.

**Additional file 10: Table of the mapped eQTL in the control, heat stress, and recovery treatment.** The eQTL are given per trait (Spot) and treatment. The QTL type, location and confidence interval is listed, as is the significance and effect. The effect is higher in N2 (positive numbers) or higher in CB4856 (negative numbers) loci. Furthermore, information about the affected gene represented by the microarray spot is shown (name, and location).

**Additional file 11: Figure of eQTL effect distribution**. (**A**) Volcano plots of the eQTL mapped per type (*cis* or *trans*) per treatment. On the x-axis the effect is plotted and on the y-axis the significance of the association is plotted (−log10(p)). Each dot represents a microarray spot and only the significant associations are shown (FDR ≤ 0.05, −log10(p) > 3.9 in all three treatments). (**B**) A histogram of the eQTL effect sizes, per type (*cis* or *trans*) per treatment. Again, the number of significantly associated spots are counted.

**Additional file 12: Table listing polymorphisms in eQTL.** The polymorphism between and CB4856 and N2 were taken from Thompson *et al.*, 2015 and counted in the genes with *cis*-, *trans*-, or no eQTL. Furthermore, the *cis*- and *trans*-eQTL have also been compared based on effect (higher expression in CB4856 or N2). The occurrences of polymorphic genes in the three sets were compared by a chi-squared test.

**Additional file 13: A list of *trans*-bands.** This table lists the identified *trans*-bands per treatment. The number of affected genes and the number of spots with a *trans*-eQTL these genes were represented by are shown.

**Additional file 14: Figure comparing allelic effects of genes with an eQTL between treatments**. A scatter plot of the effects of the eQTL of genes with an eQTL found in multiple treatments. Each dot represents a spot. The Pearson correlation values between the different comparisons are: R = 0.94 and 0.91 for *cis-* and *trans*-eQTL in control versus heat stress, R = 0.96 and 0.93 for *cis-* and *trans*-eQTL in control versus recovery, and R = 0.94 and 0.89 for *cis-* and *trans*-eQTL in heat stress versus recovery. The striped diagonal lines are shown as an optical reference.

**Additional file 15: Text detailing the calculations on overlap in *cis*- and *trans*-eQTL**.

**Additional file 16: Figure comparing genes with a *trans*-eQTL in different treatments.** (**A**). A plot of the eQTL location in treatment 1 versus the eQTL-location in treatment 2. Treatment 1 is the first treatment listed in the legend, for example: the magenta dots represent eQTL where the first treatment is control and the second treatment is heat stress. The grey lines represent the confidence interval of the eQTL based on a 1.5 drop in – log10(p). The diagonal band in this plot represents *trans*-eQTL that are regulated from the same location across treatments. (**B**) The eQTL effects of the *trans*-eQTL shown in (**A**), ordered per chromosome. The striped diagonal lines are shown as an optical reference.

**Additional file 17: A list of the enrichment analysis of genes with an eQTL**. Enrichment analysis on the genes with eQTL across different conditions. The sets were also analyzed for genes with an eQTL with a specific allelic effect, belonging to a specific *trans*-band and found, and belonging to a specific treatment. The allelic effects are indicated based on direction (*e.g. cis*-CB4856 means *cis*-eQTL higher expressed in CB4856). The database used for enrichment (Annotation) and the category (Group), and the number of genes on the array that are in the group (Genes_in_group) are indicated. Furthermore, the overlap with the cluster (Overlap) and the Bonferroni-corrected significance of that overlap are shown.

